# Toxic Effects of *Solanum sisymbriifolium* Extracts on *Meloidogyne hapla* and *Meloidogyne chitwoodi*

**DOI:** 10.64898/2026.06.25.734409

**Authors:** K. O. Chandler, H. Baker, S. W. McCotter, L. Schulz, I. Popova, H. Ibrahim, L.M. Dandurand, I. A. Zasada, C. Gleason

**Author notes:** Corresponding author: Cynthia Gleason, 1772 NE Stadium Way, Pullman, WA, 99164, USA.

## Abstract

Plant-parasitic nematodes (PPNs) are among the most destructive agricultural pests worldwide, causing significant economic losses across diverse cropping systems. Soil fumigation, the most common management strategy, is costly, detrimental to soil health, and increasingly restricted due to environmental and regulatory concerns. As a result, there is a critical need for alternative, sustainable approaches for PPN control. *Solanum sisymbriifolium* is resistant to several *Meloidogyne* species and represents a promising source of natural nematicidal compounds. In this study, we evaluated the effects of *S. sisymbriifolium* extracts on *Meloidogyne chitwoodi* and *M. hapla,* two economically important nematodes. Compounds were extracted using solvents of increasing polarity and then reconstituted in water. The water-solubilized extracts were then used in bioassays to assess their effects on nematode egg hatching, egg viability, and second-stage juvenile (J2) survival. Egg hatching and J2 viability of both *Meloidogyne* species were consistently affected by compounds in the 1-butanol fraction. Further characterization of these compounds may enable the development of novel, environmentally sustainable alternatives to conventional nematode management strategies.

---

The two most prevalent *Meloidogyne* species in the Pacific Northwest (PNW) region of the United States are *Meloidogyne chitwoodi* and *M. hapla* (Zasada et al., 2018). Both species infect a wide range of crops, including potato (*Solanum tuberosum*), which is commonly grown in the region (Zasada et al., 2018). In potato, *Meloidogyne* infections primarily reduce crop value by severely compromising tuber quality due to tuber galling and brown spots below the surface of the tuber skin (Nyczepir et al., 1982, Santo and O’Bannon, 1981). Conventional methods for the control of *M. chitwoodi* and *M. hapla* in potato have mostly relied on the use of fumigants and nematicides (Desaeger et al., 2020). Current chemical control measures are expensive, and their usage may be hazardous to human health and/or the environment. Thus, there is a need for new ways to manage nematodes that are both effective and suitable for sustainable use in agriculture (Desaeger et al., 2020).

*Solanum sisymbriifolium*, commonly known as litchi tomato or prickly nightshade, has been investigated for managing plant-parasitic nematodes in agricultural crops (Scholte, 2000, Scholte and Vos, 2000, Timmermans et al., 2007, Dandurand and Knudsen, 2016, Hajihassani et al., 2020, Perpétuo et al., 2021, Bhatta et al., 2023, Hickman and Dandurand, 2025). It has been used successfully as a *Meloidogyne*-resistant rootstock for tomato grafting (Baidya et al., 2017) and employed as a trap crop for the pale cyst nematode, *Globodera pallida*, reducing population densities by up to 99% after one year of rotation (Dandurand and Knudsen, 2016, Hickman and Dandurand, 2025, Dandurand et al., 2019). *Solanum sisymbriifolium* is also known to produce a wide range of bioactive chemicals, some of which are nematicidal (Perpétuo et al., 2023, Schulz et al., 2024). For example, *G. pallida* and *G. ellingtonae* J2 hatched in *S. sisymbriifolium* root exudates but had reduced motility and infectivity (Kud et al., 2022). Spirosolane steroidal glycoalkaloids produced by *S. sisymbriifolium*, such as α-solamargine, α-solasonine, and their aglycone, solasodine, reduced hatch of *G. pallida* (Pillai and Dandurand, 2021, Schulz et al., 2024). Meanwhile, organic acids identified in *S. sisymbriifolium* extracts (e.g. aconitic acid) immobilized *M. chitwoodi*, *M. incognita*, and *M. hapla* J2 (Jablonski et al., 2026).

In greenhouse assays, the *S. sisymbriifolium*, reduced galling on tomato inoculated with *M. arenaria*, *M. incognita*, and *M. haplanaria* compared to the control (Hajihassani et al., 2020). A further investigation using *S. sisymbriifolium* found that *M. hapla* J2 were able to penetrate and establish low levels of root infection; however, *M. chitwoodi* J2 did not enter roots and was ultimately absent from all inoculated plants (Baker et al., 2023; Kooliyottil et al., 2016).

Given that *S. sisymbriifolium* exhibits resistance to multiple nematode species such as *G. pallida* (Gómez-Armesto et al., 2025, Hickman and Dandurand, 2025), *Meloidogyne* species (Baker et al., 2023), and that most *G. pallida*, *M. hapla*, and all *M. chitwoodi* J2 are unable to penetrate the roots, we hypothesized that this resistance is mediated, at least in part, by bioactive compounds that adversely affect these nematodes (Pillai and Dandurand, 2021; Schulz et al., 2024). The goal of this study was to evaluate the toxicity of *S. sisymbriifolium* extracts prepared using a range of solvents on the eggs and J2 of *M. chitwoodi* and *M. hapla*. These findings lay the groundwork for future research exploring the biochemical basis of *S. sisymbriifolium*–mediated toxicity to *Meloidogyne* species and other plant-parasitic nematodes.

## MATERIALS AND METHODS

### Meloidogyne spp. rearing and egg collection

Populations of *M. chitwoodi* (donated by Chuck Brown, USDA-ARS, Prosser, WA) and *M. hapla* (collected from a wine grape vineyard in Veneta, OR by I. Zasada) were maintained in greenhouses (23°C, 14 hrs light:10 hrs dark) on tomato ‘Rutgers’ as described by Wram and Zasada (2020). Eggs of *M. chitwoodi* were obtained by carrying out a bulk extraction of chopped roots, while eggs of *M. hapla* were collected either by dissecting egg masses out of roots, or by bulk extraction. Eggs were cleaned via sucrose flotation according to the procedure used by Baker et al. (2023). Eggs were surface sterilized with 10% commercial bleach and thoroughly rinsed with sterile, distilled water (Gleason et al., 2017). To obtain J2, surface sterilized eggs were placed in a modified hatching chamber (Zhang and Gleason, 2021). After a 3-day incubation in the dark at room temperature, hatched J2 were collected and counted in a hemocytometer.

### Preparation of S. sisymbriifolium extracts

*Solanum sisymbriifolium* synthetic cross II (Sissyn II), used in these experiments, was developed by Dr. Chuck Brown, USDA-ARS, Prosser and seeds were obtained from Dr Dandurand at the University of Idaho. Seeds of *S. sisymbriifolium* were surface sterilized in 1% sodium hypochlorite solution and rinsed with water. Seeds were germinated in potting mix under greenhouse conditions (23°C with 14-hr light/10-hr dark photoperiod). Two weeks after planting, seedlings were transplanted into individual pots containing a 2:1 volume/volume sand-soil mixture, then grown for an additional ∼8 weeks until inflorescence. At this point they were cut at the soil line to separate stems and leaves from root tissue. The stem and leaf (SL) material was immediately flash-frozen in liquid nitrogen and subsequently lyophilized. The freeze-dried material was ground into a fine powder in a cyclone sample mill (Udy Analyzer Company, Fort Collins, CO). The processed SL material was kept in air-tight containers at −20°C until use.

A liquid-liquid extraction technique was used to extract *S. sisymbriifolium* tissues (Mazzola et al., 2008). The tissue powders, processed as described, were first extracted with methanol, then dried, and reconstituted in water (Popova and Morra, 2021). These crude extracts were further processed by using four solvents, listed here in order of increasing polarity: hexane (H), dichloromethane (D), ethyl acetate (E), and water-saturated 1butanol (B), as described by Schulz et al., 2024. The collected extract from each solvent was dried using a rotavapor R-300 (Buchi, Switzerland) and stored in methanol at −20°C. Prior to use, extracts were dried to remove solvents and reconstituted in sterile, distilled water. If needed, extracts were sonicated or vortexed for 20 seconds to help with reconstitution. Solvents are abbreviated by the first letter (H, D, E, B), and combined with stem and leaf (SL), such that stem and leaf extracts made with hexane would be abbreviated SLH.

### Preparation of 1-butanol subfractions

The five sub-fractions from the 1-butanol extracts were prepared using a solid-phase extraction technique (Ibrahim et al., 2025). In brief, 1-butanol extract was dissolved in water (100 mg/ml) and subjected to solid-phase extraction on Oasis HLB cartridges (400 mg sorbent, 60 µm; Waters, Milford, MA) preconditioned with 6 ml of methanol and 6 ml of water. The sample was loaded under 10 mm Hg negative pressure. Cartridges were dried for 10 min, washed with 5% volume/volume aqueous methanol, and eluted with 30 mL each of 10%, 30%, 50%, 80%, and 100% aqueous methanol, yielding five groups of fractions (G1 = 10%; G2 = 30%; G3 = 50%, G4 = 80%; and G5 = 100%). Fractions were stored in methanol in a −20°C freezer until use, at which point they were resuspended in water.

### *Meloidogyne* species microwell assays with extracts

A series of experiments were conducted to determine effects of extracts on *M. chitwoodi* and *M. hapla.* All experiments were conducted in transparent 96-well, flat-bottomed culture plates. Each well received 190 µl of its respective water-resuspended *S. sisymbriifolium* extract (SLD, SLH, SLE, or SLB) and 10 µl of aqueous nematode suspension. For assays with *Meloidogyne* J2, 30 to 60 J2 were added per well. For assays with *Meloidogyne* eggs, 100 to 300 eggs were added per well. Sterile, distilled water was used as a control. The plates were incubated at room temperature in the dark for 24 hours for J2 assays and 7 days for egg assays. Second-stage juveniles were identified as living based upon motility upon agitation or when 4 µl 1 N NaOH was added to the well (Chen and Dixon, 2000). Egg hatch was determined by counting number of eggs at day 0 and day 7.

An additional experiment investigating hatching of *M. chitwoodi* eggs was performed using a five-part dilution of the SLE and SLB extracts. The dilutions were made by mixing 1 part extract with 4 parts water/egg solution. The percentage egg hatch was measured after 5 and 14 days of exposure. After the 14-day incubation, any unhatched eggs in each well were rinsed then resuspended in 1 mL of water, then mixed with 1 mL of Meldola’s Blue stain (0.5% weight/volume). After 48 hours, the eggs were rinsed with sterile distilled water. Dead eggs absorbed the stain and turned dark purple while live eggs remained clear.

In all experiments, the egg hatch percentage was calculated as (number of J2 hatched/number of eggs at time of inoculation) * 100 = percentage hatch. Second-stage juvenile percentage mortality was calculated as (number of dead J2 after 24 h /number of initial J2 in each well) * 100 = percentage mortality. In all hatching assays, treatments were replicated at least five times, and each experiment was repeated at least twice for each species of nematode. In the dilution experiment, egg viability percentage was calculated using (live eggs/total eggs) * 100 = percent viability. Egg viability experiments were repeated twice with treatments replicated 7 - 8 times.

### Meloidogyne species greenhouse experiments with extracts

Experiments were conducted with *M. hapla* and *M. chitwoodi* to assess effects of extractions on invasion. For *M. hapla,* tomato ‘Rutgers’ seeds were planted as described above for *S. sisymbriifolium*. *M. hapla* eggs (1,500 eggs) were placed in 1.5 mL tubes containing either undiluted extracts (SLE, SLD, SLB, or SLH) or water (control) and incubated in the dark at room temperature for 1 or 3 days. After incubation, tubes were lightly vortexed, and 500 µL of water was added to dilute the suspensions. Prior to inoculation, tomato roots were trimmed to approximately equal lengths, and plants were transplanted into approximately 100 g of a 1:1 soil/loam mixture in 6-pack plastic containers. Inoculation was performed by making two holes approximately 3 cm deep in the substrate using a 1 mL pipette tip. The holes were positioned on opposite sides of the plant, approximately 1 cm from the stem. Approximately 250 eggs exposed to the extracts were dispensed in two 500 µL aliquots, one into each hole. The holes were then covered, and plants were maintained in the greenhouse for the duration of the experiment. Tomato plants were evaluated seven weeks after inoculation. Shoots were removed at the soil line, and egg masses were extracted from the roots using 1% sodium hypochlorite (Gleason et al., 2017). Roots were thoroughly rinsed to remove bleach and dried in a 70°C oven for seven days before determining dry root weight. Eggs were counted using an inverted microscope, and data were expressed as reproduction factor (RF = final egg density/initial egg density). The experiment was repeated at least twice.

For *M. chitwoodi,* tomato *‘*Rutgers’ seeds were planted in the same fashion as described for *S. sisymbriifolium*. Two weeks after germination, the tomato plants were transplanted into cone-tainers containing ∼100 g of sand and grown in the greenhouse for two more weeks before inoculation. *M. chitwoodi* eggs (9,000 eggs) were placed in either five-part dilutions of SLB or SLE (1 part extract: 4 parts water with eggs) or sterile distilled water (control) and incubated in the dark at room temperature for five days. Each tomato plant was then inoculated with approximately 1,000 eggs by making two holes approximately 3 cm deep in the substrate using a 1 mL pipette tip. The holes were positioned on opposite sides of the plant, approximately 1 cm from the stem. Eggs in the treatments were dispensed in two 500 µL aliquots, one into each hole. The holes were then covered, and plants were maintained in the greenhouse for the duration of the experiment. Tomato plants were evaluated for root galling at four weeks post-inoculation. The plants were cut at the soil level to remove above-ground tissue. Roots were rinsed twice in water to remove the sand, dried, and then weighed. To help visualize galls, the roots were stained for five minutes with a 1% Ponceau stain and rinsed with water. Galling was evaluated under a stereo microscope (Stemi 2000-C, Zeiss, Germany). Severity of galling was measured by calculating galls per gram of root. The experiments were conducted twice.

### Data analysis and normalization

Data analyses and figure preparation were conducted using GraphPad Prism 10.6.1 (GraphPad Software, Boston, MA). Experiments were conducted twice. Prior to pooling the data, differences among trials were assessed using a one-way ANOVA followed by Tukey’s multiple-comparisons test. When no significant differences were detected among trials (P > 0.05), data from all trials were combined for subsequent analyses. Data that were not normally distributed following post hoc assessment were analyzed using the Mann–Whitney test, whereas normally distributed data were analyzed using Welch’s *t*-test, where indicated.

## RESULTS

### Meloidogyne hapla pre-treated with S. sisymbriifolium stem and leaf extracts affected J2 viability, egg hatching, and reproduction

The effects of four *S. sisymbriifolium* extracts (SLB, SLH, SLD, and SLE) on *M. hapla* J2 were evaluated after 24 hours of incubation (Fig. 1). The water control exhibited the lowest mean percentage of dead *M. hapla* J2 (0.2%), confirming the reliability of the mobility assay. Among the treatments, SLB resulted in the highest *M. hapla* J2 mortality at 24 hours, 48%. Overall, all extracts demonstrated some degree of toxicity toward *M. hapla* J2. To determine the effects of the extracts on egg hatching, *M. hapla* eggs were incubated for 7 days in water or the *S. sisymbriifolium* extracts (Fig. 2). All extracts resulted in statistically significant reductions in *M. hapla* egg hatching relative to the water control. The greatest average reduction in hatching was observed with SLB, followed by SLE.

**Figure 1.**
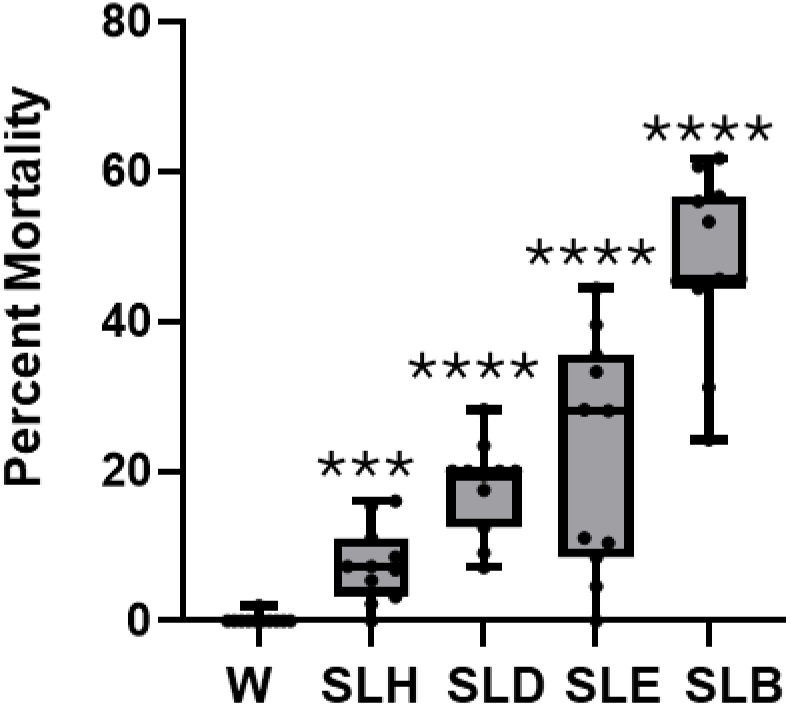
Exposure to *Solanum sisymbriifolium* extracts for 24 h increased *Meloidogyne hapla* J2 mortality compared to a water control. Box plots show the median percentage of dead J2 for each treatment. Box limits represent the 25th and 75th percentiles, and whiskers indicate the minimum and maximum values. W, water treatment; SLH, stem and leaf hexane; SLD, stem and leaf dichloromethane; SLE, stem and leaf ethyl acetate; SLB, stem and leaf 1-butanol. Data shown are two independent experiments combined. (N = 10). Statistical significance compared to water control was determined using the Mann–Whitney test (*** P < 0.005, ****P < 0.0005). Treatments are presented in order of increasing solvent polarity used for extraction.

**Figure 2.**
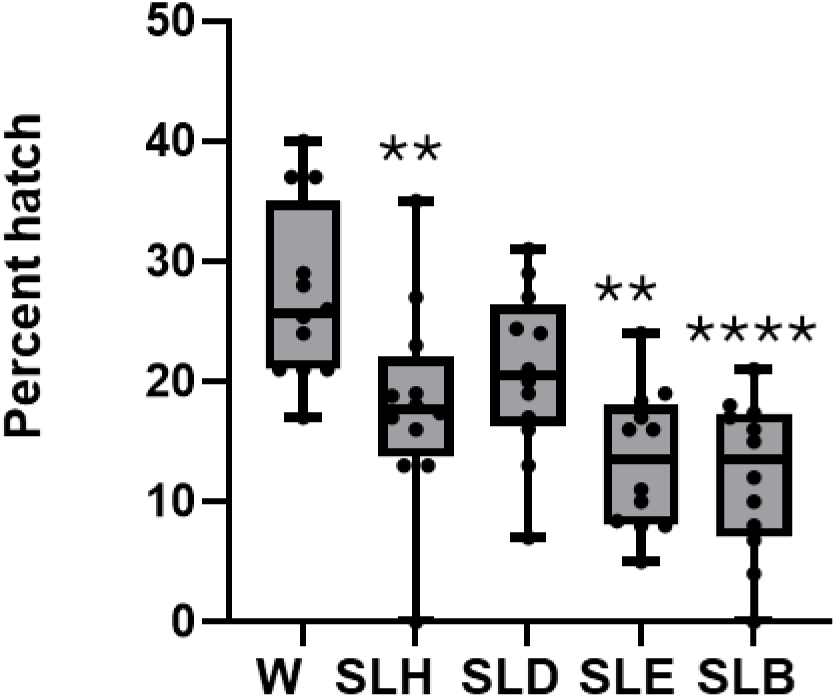
Exposure to *Solanum sisymbriifolium* extracts for 7 days decreased *Meloidogyne hapla* egg hatching compared to a water control. Box plots show the median percentage of egg hatching for each treatment, relative to water (W) set to 100%. Box limits represent the 25th and 75th percentiles, and whiskers indicate the minimum and maximum values. W, water treatment; SLH, stem and leaf hexane; SLD, stem and leaf dichloromethane; SLE, stem and leaf ethyl acetate; SLB, stem and leaf 1-butanol. Data shown are two independent experiments combined. (N = 12). Statistical significance was determined using the Mann–Whitney test (\*\**P* < 0.005, \*\*\*\**P* < 0.0005). Treatments are presented in order of increasing solvent polarity used for extraction.

Next, to assess if a pretreatment of *M. hapla* eggs with the extracts (SLB, SLH, SLD, and SLE) would affect their ability to infect and reproduce on a host plant, the eggs were incubated in the extracts for either 24 or 72 h, rinsed with water, and then used to inoculate susceptible tomato plants. *M. hapla* eggs exposed to extracts for 24 h did not exhibit significant differences in RF values compared to the water control (Fig. 3A). In contrast, eggs incubated in SLB for 24 h had a slight increase in RF. Similarly, *M. hapla* eggs exposed to SLB and SLH extracts for 72 h exhibited slight increases in RF values relative to the control (Fig. 3B).

**Figure 3.**
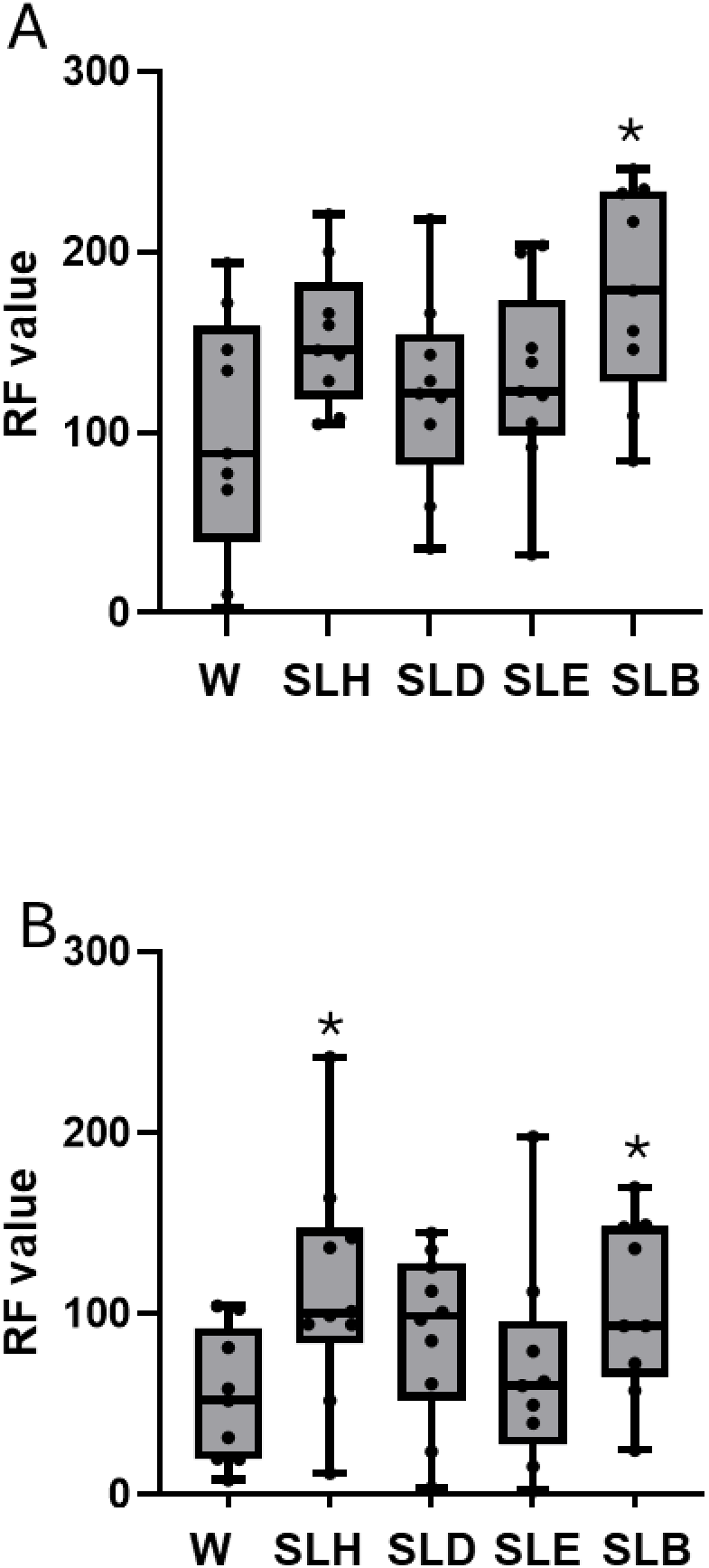
Exposure of *Meloidogyne hapla* eggs to *Solanum sisymbriifolium* extracts for (A) 24 h and (B) 72 h affected reproduction factor (RF = final density/initial density) values. Box plots show the median RF value for each treatment. Box limits represent the 25th and 75th percentiles, and whiskers indicate the minimum and maximum values. W, water treatment; SLH, stem and leaf hexane; SLD, stem and leaf dichloromethane; SLE, stem and leaf ethyl acetate; and SLB, stem and leaf 1-butanol. Data from two independent experiments were combined (N = 10). Statistical significance for panels A and B was determined using the Mann–Whitney test compared with the water control (*P < 0.05). Treatments are presented in order of increasing solvent polarity used for extraction.

### S. sisymbriifolium stem and leaf extracts reduced M. chitwoodi J2 viability and egg hatching

Because *S. sisymbriifolium* stem and leaf extracts significantly increased *M. hapla* J2 mortality after 24 h of incubation (Fig. 1A), we next evaluated whether these extracts had similar effects on *M. chitwoodi* J2. Similar to the effect seen for *M. hapla* J2, all extracts resulted in a significant increase in *M. chitwoodi* J2 mortality (Fig. 4). The SLB extract had the strongest effect on the *M. chitwoodi* J2, causing approximately 21% mortality. *Meloidogyne chitwoodi* egg hatching was also influenced by the *S. sisymbriifolium* extracts (SLB, SLH, SLD, or SLE) after seven days of exposure (Fig. 5). Exposure to SLB and SLE significantly reduced *M. chitwoodi* egg hatch, whereas SLH increased hatching compared to the water control. In contrast, SLD had no significant effect on *M. chitwoodi* egg hatch.

**Figure 4.**
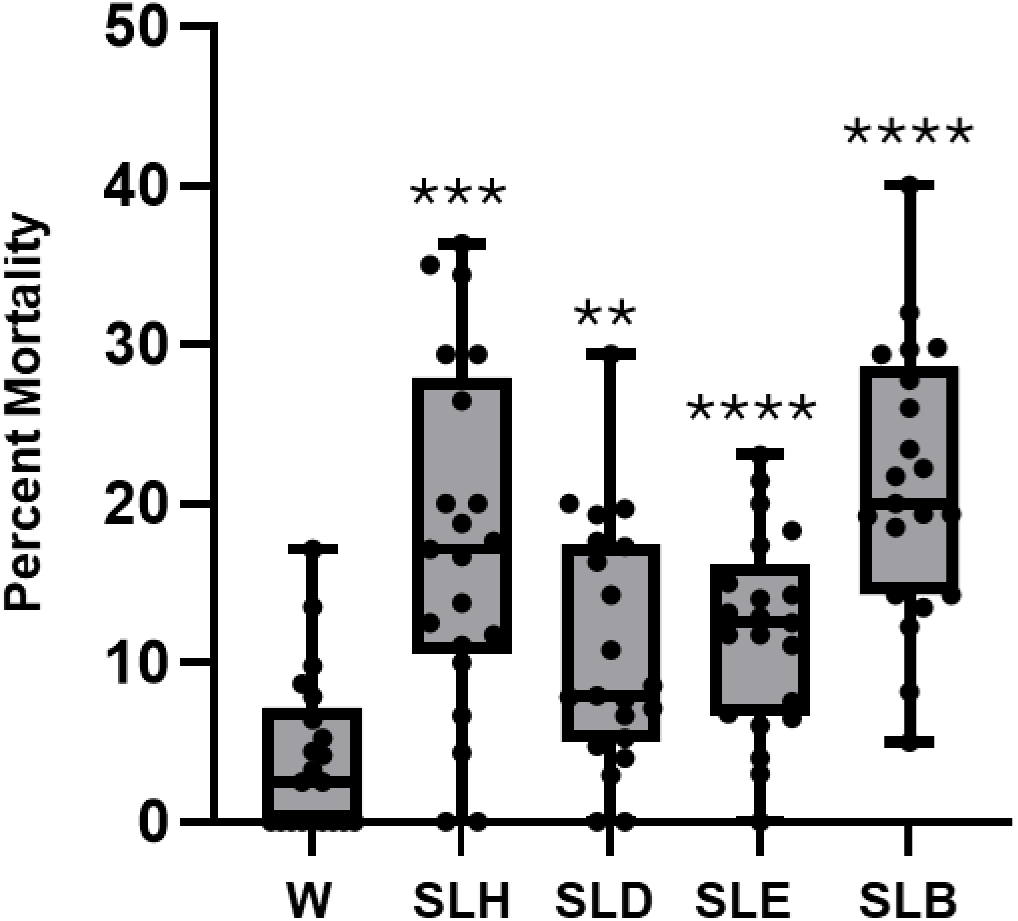
Exposure to *Solanum sisymbriifolium* extracts for 24-h affected *Meloidogyne chitwoodi* J2 mortality. Box plots show the median percentage of dead J2 after incubation in water (W; control) or the following extracts: SLH, stem and leaf hexane; SLD, stem and leaf dichloromethane; SLE, stem and leaf ethyl acetate; and SLB, stem and leaf 1-butanol. Box limits indicate the 25th and 75th percentiles, and whiskers represent the minimum and maximum values. Data from two independent experiments were combined (N = 21). Statistical significance was determined using the Mann–Whitney test compared with the water control (**P < 0.005; ***P < 0.001; ****P < 0.0005).

**Figure 5.**
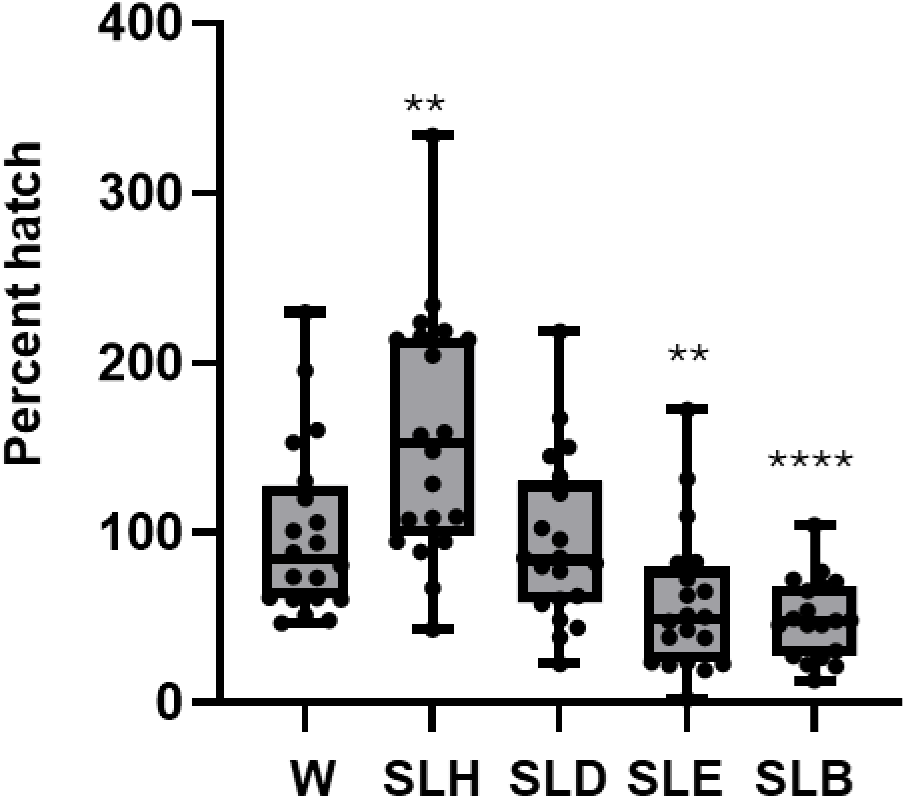
Exposure of *Meloidogyne chitwoodi* eggs to *Solanum sisymbriifolium* extracts for 7 days affected egg hatching. Box plots show the median percentage of hatched eggs after incubation in the extracts relative to hatch in the water control. Extracts included SLH, stem and leaf hexane; SLD, stem and leaf dichloromethane; SLE, stem and leaf ethyl acetate; and SLB, stem and leaf 1-butanol. Box limits represent the 25th and 75th percentiles, and whiskers indicate the minimum and maximum values. Data from two independent experiments were combined (N = 20–21). Statistical significance relative to the water control was determined using the Mann–Whitney test (**P < 0.005; ****P < 0.0005).

Because the SLB extract had the most consistent and toxic effects on *M. chitwoodi* egg hatching and J2 viability, it was sub-fractionated into five (1-5) different fractions, numbered according to increasing polarity. Next, we assessed whether these subfractions had effects on *M. chitwoodi* egg hatching and J2 viability. All subfractions (SLB1, SLB2, SLB3, SLB4, and SLB5) significantly reduced *M. chitwoodi* egg hatching relative to the water control (Fig. 6). Subfractions SLB1 and SLB2 caused the lowest median percentages of egg hatching, while SLB3, SLB4, and SLB5 also produced statistically significant reductions compared with the control (Fig. 6).

**Figure 6.**
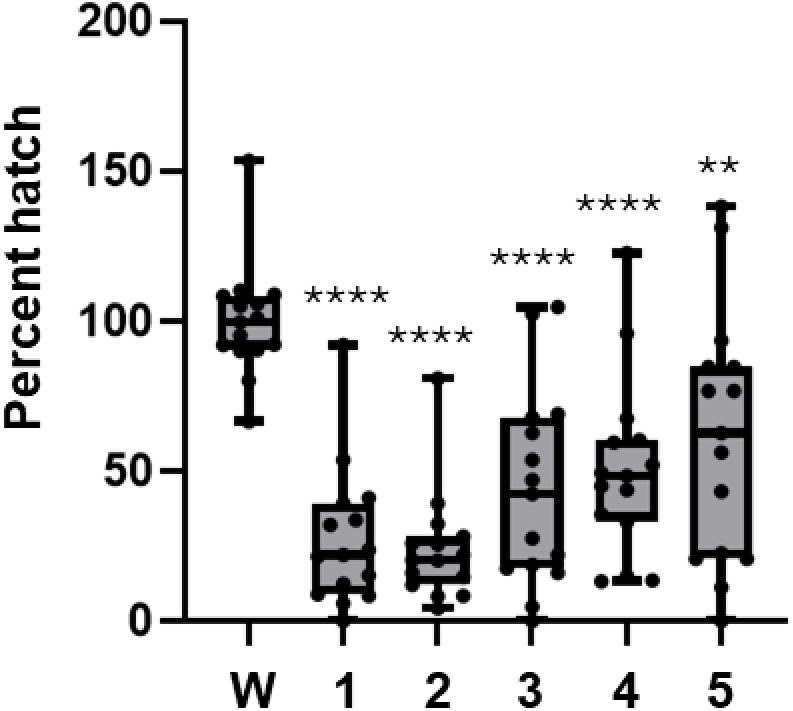
Exposure of *Meloidogyne chitwoodi* eggs to *Solanum sisymbriifolium* 1-butanol (SLB) subfractions 1–5 for 7 days affected egg hatching. Box plots show the median percentage of egg hatching for each treatment relative to hatch in the water control (W). Box limits represent the 25th and 75th percentiles, and whiskers indicate the minimum and maximum values. Treatments are presented in order of increasing solvent polarity used for extraction. Data from two independent experiments were combined (N = 20). Statistical significance relative to the water control was determined using the Mann–Whitney test (**P < 0.005; ****P < 0.0005).

### Butanol and ethyl acetate extracts were effective in killing M. chitwoodi eggs and reducing host plant infections

Due to limited quantities of material, we focused on *M. chitwoodi* as a representative *Meloidogyne* for subsequent assays and employed five-part (1:4) dilutions of the SLB and SLE extracts. Incubating *M. chitwoodi* eggs in the SLB and SLE extracts for 5 days did not reduce egg hatch relative to the water control (Fig. 7A). However, after a 14-day incubation, both extracts significantly suppressed *M. chitwoodi* egg hatching compared with the control (Fig. 7A). Assessment of viability using Meldola Blue staining of remaining unhatched eggs after 14-day incubation in extracts revealed that most *M. chitwoodi* eggs were dead following exposure to five-part (1:4) dilutions of the SLB and SLE extracts (Fig. 7B). When eggs exposed for 5 days to SLB and SLE were used to inoculate tomato plants, pretreatment with SLE had no significant effect on root galling, whereas SLB reduced galling by an average of 40% (Fig. 8).

**Figure 7.**
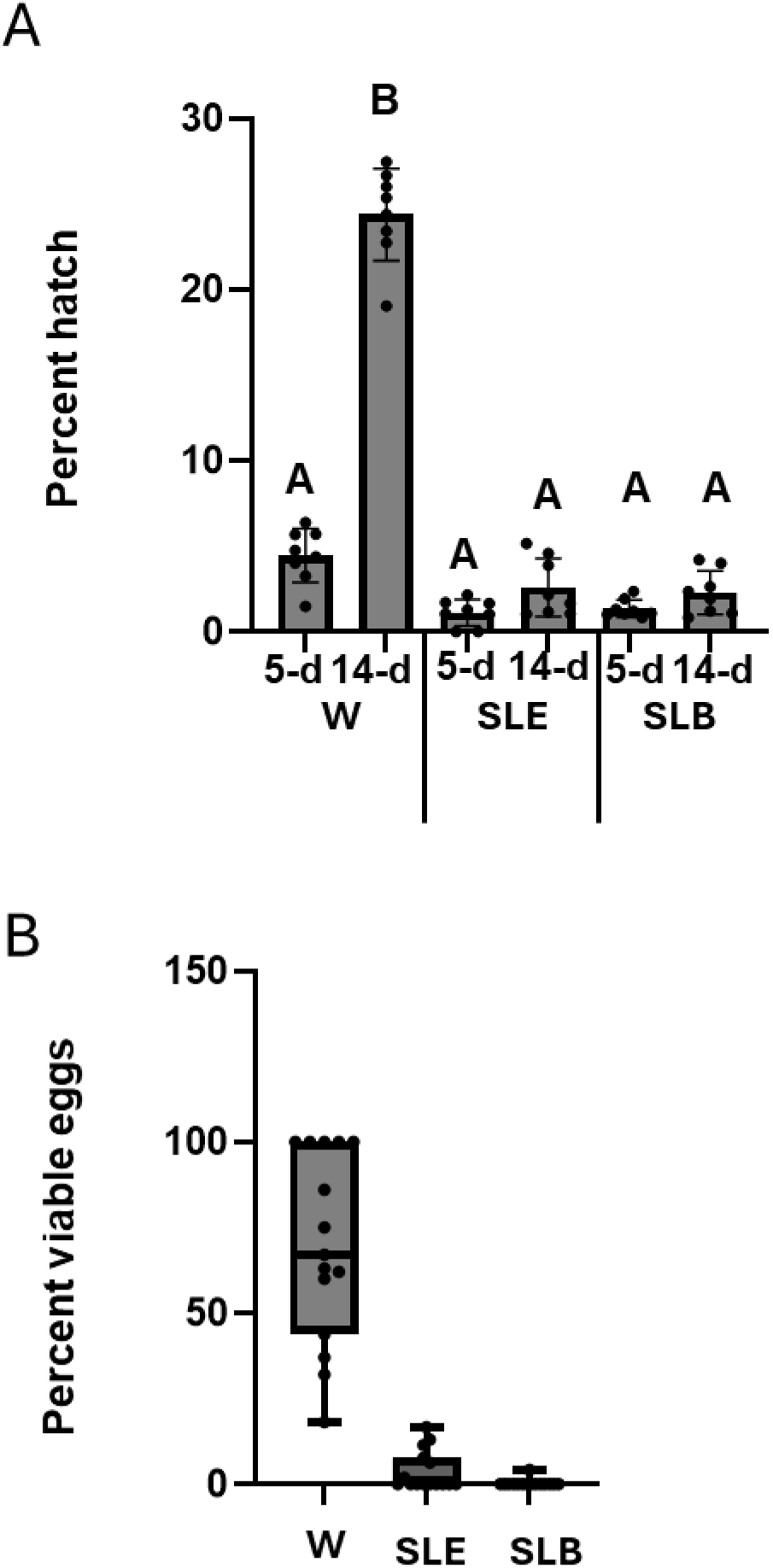
Exposure to *Solanum sisymbriifolium* 1-butanol (SLB) and ethyl acetate (SLE) diluted (1:4) extracts for 14 days affects *Meloidogyne chitwoodi* egg hatching and egg viability. (A) Bar graphs show mean hatch percentage after 5-days (d) and 14-d in water (W) control or in SLE and SLB treatments +/- standard deviation (SD), (N = 7). Experiment repeated with similar results. Different letters indicate significant differences determine by Tukey HSD. B) Box plots show median percentage of egg viability after 14-d incubation in water, SLE or SLB extracts +/-SD as determined with Medola blue staining. (N=15). Statistical significance relative to the water control was determined using the Mann–Whitney test (****P <0.0001).

**Figure 8.**
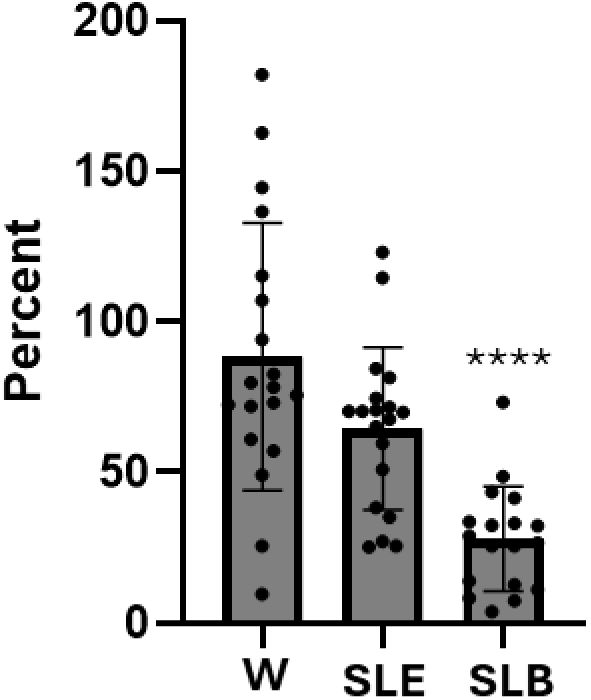
Exposure of *Meloidogyne chitwoodi* eggs to *Solanum sisymbriifolium* 1-butanol (SLB) and ethyl acetate (SLE) extracts reduced infection of tomato plants. Bar graphs show the mean number of galls per gram of root relative to the water control (W) ± SD. Data from two independent experiments were combined (N = 18-19). Statistical significance was determined using Welch’s *t*-test compared with the water control (****P < 0.0005).

## DISCUSSION

Previously, Schulz et al. (2024) reported that extracts from *S. sisymbriifolium* reduced egg hatch and viability of *G. pallida*, with the SLB and SLH extracts having particularly strong effects, reducing egg hatch 49.5%, and 68.3%, respectively. Building on these findings, the objective of the present study was to determine whether extracts generated using the same solvents also exhibit toxicity toward *M. hapla* and *M. chitwoodi*. In contrast to what was observed for *G. pallida*, all stem and leaf extracts tested in this study exhibited some degree of toxicity toward the *Meloidogyne* spp. For *M. hapla*, the SLB extract consistently resulted in the greatest J2 mortality. Exposure of *M. hapla* eggs to the extracts for seven days significantly reduced egg hatching, indicating that these extracts all have compounds that are toxic to *M. hapla* eggs. To further examine potential shorter-term effects, *M. hapla* eggs were exposed to the extracts for 24 hours and then used to inoculate susceptible tomato plants. A 24-hour exposure to SLH, SLD, or SLE had no effect on *M. hapla* RF values, suggesting that this duration was insufficient to impair egg hatch or subsequent infection processes. However, a 24-hour exposure to SLB resulted in a slight increase in *M. hapla* RF values. Similar increases were observed following 72-hour exposures to SLB and SLH. For *M. chitwoodi*, the SLB extract consistently had the greatest effect on J2 mortality, resembling the effects observed for *M. hapla.* In terms of egg hatching, the SLB, SLE, and SLH extracts had the most pronounced effects on *M. chitwoodi* egg hatching after a 7-day exposure. While SLB and SLE produced the greatest reductions in egg hatch, SLH was associated with increased hatching.

Although egg hatching assays are inherently variable, this variability was particularly evident in the effects of the undiluted SLH fraction on both *M. hapla* and *M. chitwoodi* egg hatching. Such inconsistency may indicate that the hexane-derived fraction contains a variable or unstable mixture of bioactive compounds, leading to less predictable biological activity. The increased *M. chitwoodi* hatching observed after seven days of exposure to the SLH fraction suggests that prolonged treatment with this extract may stimulate egg hatch rather than suppress it. The elevated RF values observed for *M. hapla* eggs treated for 72 hours with SLH further support this possibility.

Because pretreatment of *M. chitwoodi* eggs with SLB and SLE reduced both egg viability, hatch, and J2 survival, these extracts were subsequently used to treat eggs prior to plant inoculation. In these experiments, root galling was significantly reduced after SLB treatment compared to the water control. Together, these findings suggest that the SLB extract contains bioactive compounds capable of negatively affecting *M. chitwoodi* and reducing plant infection.

For both *Meloidogyne* species, the SLB extract was the most consistently effective at reducing egg hatching, mirroring previous findings in *G. pallida* (Schulz et al., 2024). Therefore, the SLB extracts were further fractionated based on water solubility into five fractions to identify active compound(s) that may be contributing to toxicity against *Meloidogyne*. Based on solvent extraction, fraction 1 contained the least polar compounds and fraction 5 contained the most polar. The first two fractions were more toxic to *M. chitwoodi* eggs compared to the last 3 fractions. We predicted that the more polar fractions would contain higher levels of glycoalkaloids based on predicted solubility. Glycoalkaloids are toxic to plant-parasitic nematodes (Perpétuo et al., 2023; Schulz et al., 2024). Specifically, the glycoalkaloid α-solamargine, which was concentrated in SLB extracts, was toxic to *G. pallida* (Schulz et al., 2024). The sub-fraction that likely contained these glycoalkaloids, based on solubility, was the least effective against *M. chitwoodi*, whereas other fractions showed greater activity. This suggests that compounds other than glycoalkaloids may contribute to toxicity against *M. chitwoodi*. It is possible that *M. chitwoodi* may be sensitive to a broader or different range of metabolites compared to *G. pallida*.

While SLE extracts had no impact on *G. pallida*, they were found to be toxic to *M. chitwoodi* and *M. hapla*, again suggesting differences in species-specific sensitivities. Ethyl acetate is known to extract phenolic acids and flavonoids (Gopčević et al., 2022) and certain flavanols such as kaempferol, quercetin, and myricetin suppressed other plant-parasitic nematodes (Wuyts et al., 2006). These compounds may play a role in the observed toxicity, although metabolite identification and further sub-fractionation are needed to confirm this.

Overall, these findings underscore the potential of *S. sisymbriifolium* metabolites, particularly those extracted with ethyl acetate and 1-butanol, as sources of novel nematicidal compounds. Continued characterization of these extracts and their active fractions may facilitate the development of targeted, environmentally friendly tools for plant-parasitic nematode management.

## Acknowledgements

Support was provided by the U.S. Department of Agriculture-National Institute of Food and Agriculture SCI CA grant 2022-51181-38450. CG was supported in part by USDA NIFA Hatch WNP7003632 and Multistate Hatch Project W5186.

## Notes

### Competing Interest Statement

The authors have declared no competing interest.

